# The molecular mechanism of the type IVa pilus motors

**DOI:** 10.1101/082750

**Authors:** Matthew McCallum, Stephanie Tammam, Ahmad Khan, Lori L. Burrows, P. Lynne Howell

## Abstract

Type IVa pili are protein filaments essential for virulence in many bacterial pathogens; they extend and retract from the surface of bacterial cells to pull the bacteria forward with unprecedented force. They are used for attachment, swarming and twitching motility, biofilm formation, up-regulation of other virulence factors, and natural competence. The pilus is assembled by the motor subcomplex which consists of the inner membrane protein PilC and the cytoplasmic ATPase PilB. How PilB catalyzes this process is unknown, due in part to the lack of high-resolution structural information. Phylogenetic analysis of PilB-like ATPases, including GspE, PilT, BfpD, FlaI, and archaeal GspE2 revealed highly conserved residues essential for function in this family of ATPases. Here we report the structure of the core ATPase domains of *Geobacter metalloreducens* PilB bound to ADP and the non-hydrolysable ATP analogue, AMPPNP, at 3.4 and 2.3Å, respectively. Importantly, these structures were determined in non-saturating nucleotide conditions, revealing important differences in nucleotide binding between chains. Analysis of these differences revealed the sequential turnover of nucleotide by the chains, and the corresponding domain movements. Our data indicate a clockwise rotation of movement in PilB, which would support the assembly of a right-handed helical pilus. Conversely, our analysis suggests a counterclockwise rotation in PilT that would enable right-handed pilus disassembly. The proposed model provides insight into how this family of ATPases can power pilus extension and retraction with extraordinary forces.

## INTRODUCTION

The Type IVa Pilus (T4aP) is a protein grappling hook that can pull bacteria forward with impressive forces that can exceed 100 pN [1]. The bacteria extend these pili to attach to surfaces, and retract them to pull the bacteria towards the point of attachment, mediating irreversible attachment or surface associated twitching motility [2]. The T4aP is homologous to the type IVb pilus, the type II secretion system, archaeal flagella, and bacterial competence systems [3, 4]. Collectively, these machines can be identified in every major phylum of prokaryotic life [5].

Despite the importance of the T4aP and homologous systems, little is known about how the motors of these machines work. It is thought that the energy is provided by hexameric ATPases in the cytoplasm, known as PilT-like ATPases. Well-characterized examples of PilT-like ATPases include PilB and PilT of the T4aP, GspE of the type II secretion system, FlaI of the archaeal flagella, and VirB11 of the type IV secretion system [6]. PilT-like ATPases are a family within the Additional Strand Catalytic “E” (ASCE) superfamily of ATPases, and as such are related to—but phylogenetically distinct from—FtsK-like ATPases and AAA+ ATPases [7]. Enzymes in the ASCE superfamily contain a Walker A motif, GXXXXGK[ST], and Walker B motif, hhhh[DE], (where X is any residue and h is a hydrophobic residue) used for binding the phosphates of ATP and coordinating a magnesium ion, respectively [8]. The Walker B motif of PilT-like ATPases is atypical in that the acidic residue essential for magnesium coordination is replaced with glycine [9]. Immediately following the Walker B motif is a glutamate involved in coordinating water for hydrolysis of the ɣ-phosphate of ATP [10]. PilT-like ATPases also contain conserved histidines in a unique HIS-box motif, and conserved acidic residues in a unique ASP-box motif [11]. While mutations to these motifs in PilT-like ATPases disrupt ATPase activity and *in vivo* function [9, 11, 12], the specific functions of the Walker B, HIS-box, and ASP-box motifs in PilT-like ATPases are not understood, providing an incomplete picture for how ATP hydrolysis may power T4aP-like systems.

The T4aP system has two PilT-like ATPases: PilB and PilT. PilB is thought to promote the multimerization of PilA monomers into a long helical polymer known as the pilus; this polymerization leads to pilus extension [2]. Conversely, PilT is thought to facilitate pilus retraction by depolymerizing the PilA polymer [13]. The manner in which PilB or PilT control PilA polymerization is a mystery, as the only cytoplasmic region of PilA is a short leader sequence that is cleaved at the inner face of the cytoplasmic membrane prior to polymerization [14]. Pull-down experiments indicated that PilB interacts with the N-terminal domain of PilC (PilC^NTD^) leading us to predict that PilC might bridge the gap between PilA and PilB/PilT by binding PilA in the inner membrane and PilB/PilT in the cytoplasm [15]. This prediction is consistent with the recent cryo-electron tomography-derived model of the T4aP machinery [16]. In this study it was hypothesised that PilB and PilT might function by rotating PilC to stimulate PilA polymerization or depolymerisation [16].

The currently available structures of PilT-like ATPases do not shed light on how these enzymes could turn PilC. This is in part related to the heterogeneity of the PilT-like ATPase crystal structures. PilT and GspE were crystallized as circular C6 symmetric hexamers, while PilT, GspE, and FlaI crystallized as oblong C2 symmetric hexamers, and FlaI and archaeal GspE2 as C3 symmetric hexamers [12, 17-20]. While it is not clear which conformation is more representative of PilT-like ATPases *in vivo*, it is worth noting that a C6 hexamer would only represent a functional state if all six chains simultaneously bound and catalyzed ATP, as proposed for the SV40 large T-antigen [21]. It is difficult to envision how this could rotate PilC. In contrast, most ASCE ATPases are thought to use a rotary mechanism for ATP turnover operating with either no symmetry, C2 symmetry, or C3 symmetry [10]. The C2 symmetric structures of PilT and GspE, and the C3 symmetric structure of FlaI have failed to decisively suggest a model for ATP binding and turnover because the resolution or nucleotide occupancy of these structures was not sufficiently high enough to unambiguously identify bound nucleotides [17, 19, 20]. The C2 symmetric structure of FlaI and the C3 symmetric structure of archaeal GspE2 are of sufficient resolution to identify bound nucleotides, but during crystallization these proteins were saturated with ATP or AMP-PNP, respectively, and as a result all six sites are occupied by the same nucleotide [12, 20]. With all sites occupied by the same nucleotide, it is difficult to conclude with certainty which sites in a hexamer have high/low affinity for ATP and/or ADP.

Since there is no high-resolution structural information on PilB, a major sub-family of PilT-like ATPases, we determined the structure of PilB. Using limiting concentrations of nucleotide during PilB crystallization, we were able to provide strong evidence for the relative affinities of nucleotides for various sites. With this information, we have developed a model for ATP binding and turnover in PilB that suggests a general mechanism of PilT-like ATPase function.

## RESULTS

### Phylogenetic analysis highlights highly conserved residues

To organize the members of the PilT-like ATPases into sub-families, the sequences of PilB-like ATPases were aligned and a phylogenetic tree was created as described in the Experimental Procedures (FIGURE 1A). As expected, proteins with similar functions clustered together. PilT and PilU are tightly clustered, and cluster near the BfpF retraction ATPase. PilB and GspE also clustered together. BfpD, TcpT, PilQ, CofH, LngH extension ATPases from the type IVb pilus clustered together. Archaeal GspE2 clustered closely with FlaI. These results are similar to those obtained previously [6].

**Figure 1.**
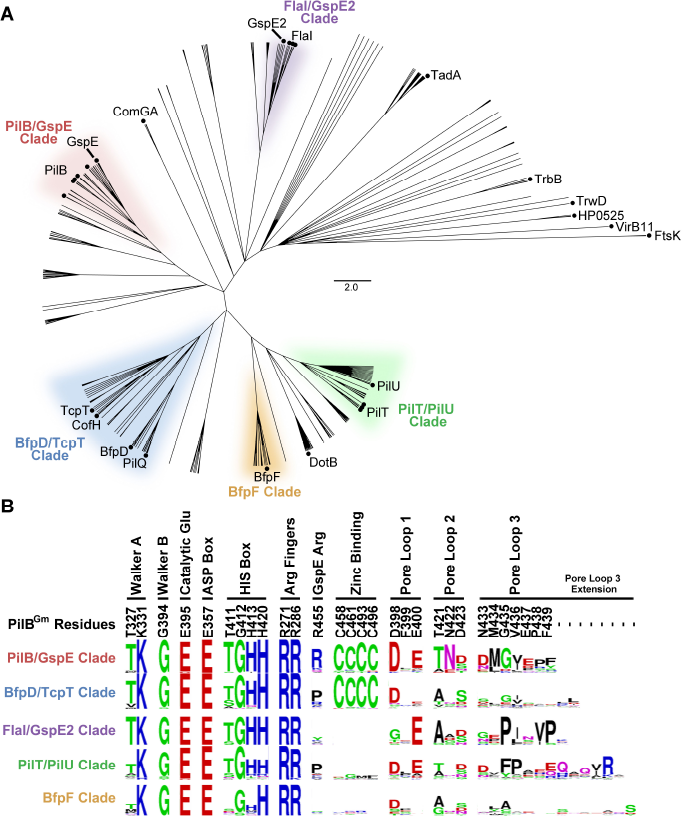
Identification and function prediction of conserved residues in PilB. **(A)** Overview of the phylogenetic analysis of PilT-like ATPase family members. Sequences from model systems are identified with a black circle and labeled. There are multiple circles for PilB, PilT, and FlaI reflecting that there are multiple model systems from different species for these proteins. For a detailed view of the phylogenetic tree including the identity of branches not labeled here see Supplemental Figure 1. Only branches with a >85% bootstrap value are shown (1000 bootstraps). FtsK was used as an out-group. Protein sub-families, labeled as clades, are given a unique color for clarity and to stratify the sequences for further analysis. **(B)** Sequence logo representation of conserved residues stratified based on the clade definitions above. Each dash indicates that there is no corresponding residue in PilB from *G. metalloreducens*.

The conservation of important residues in each sub-family or clade was plotted as sequence logos, revealing residues with complete conservation in this family (FIGURE 1B and Supplemental Figure 1): residues in the Walker A and B motifs, the catalytic glutamine, two arginine fingers, and a glutamate of the ASP box. A histidine of the HIS box was also highly conserved. We also identified residues conserved within sub-families, but not between sub-families: for instance, a tetra-cysteine zinc-binding motif, characterized in GspE and PilB [22, 23], was conserved the PilB/GspE clade as well as the BfpD/TcpT clade. Interestingly, an arginine previously shown to mediate an inter-chain contact in hexameric GspE [19] was conserved throughout the PilB/GspE clade.

### PilB is an Elongated Hexamer with C2 symmetry

The phylogenetic clustering of GspE with PilB, as well as the conservation of an arginine used to mediate an inter-chain contact, suggested that PilB may form a hexamer similar to that of GspE (PDB 4KSR). After screening several PilB constructs, we found that PilB from *G. metalloreducens* formed crystals that diffracted anisotropically to 3.4Å in one dimension and 3.9Å in the other two dimensions. Screening of many different crystals and crystal conditions revealed the anisotropy to be a consistent phenomenon. The structure of PilB was solved by molecular replacement using GspE as the search model and we were able to build residues 181 to 568 in each chain. These residues encompass the second N-terminal domain (N2D), and the C-terminal domain (CTD), residues 181-288 and 296-568, respectively. A flexible linker, residues 289-295, connects the two domains. Despite starting with full-length PilB, the first N-terminal domain (N1D) of PilB could not be built into the electron density. Removing these residues from the construct did not disrupt crystallization or improve diffraction quality.

Three chains could be built in the asymmetric unit, and an elongated hexamer was identified in the crystal packing (FIGURE 2A). Consistent with our phylogenetic analysis, the PilB hexamer was most similar to the elongated GspE hexamer (RMSD^Cα^ 2.7Å / chain, RMSDCα 5.9Å / entire hexamer). There is anomalous signal in the predicted tetra-cysteine zinc-binding sites, so zinc was also modelled into these motifs in each chain. There is density in the predicted ATP binding sites of all six chains. In four sites, ADP and magnesium best fit the electron density. For two sites, the density was too small for ADP. Instead, formate, which was present in the crystallization buffer, was modelled.

**Figure 2.**
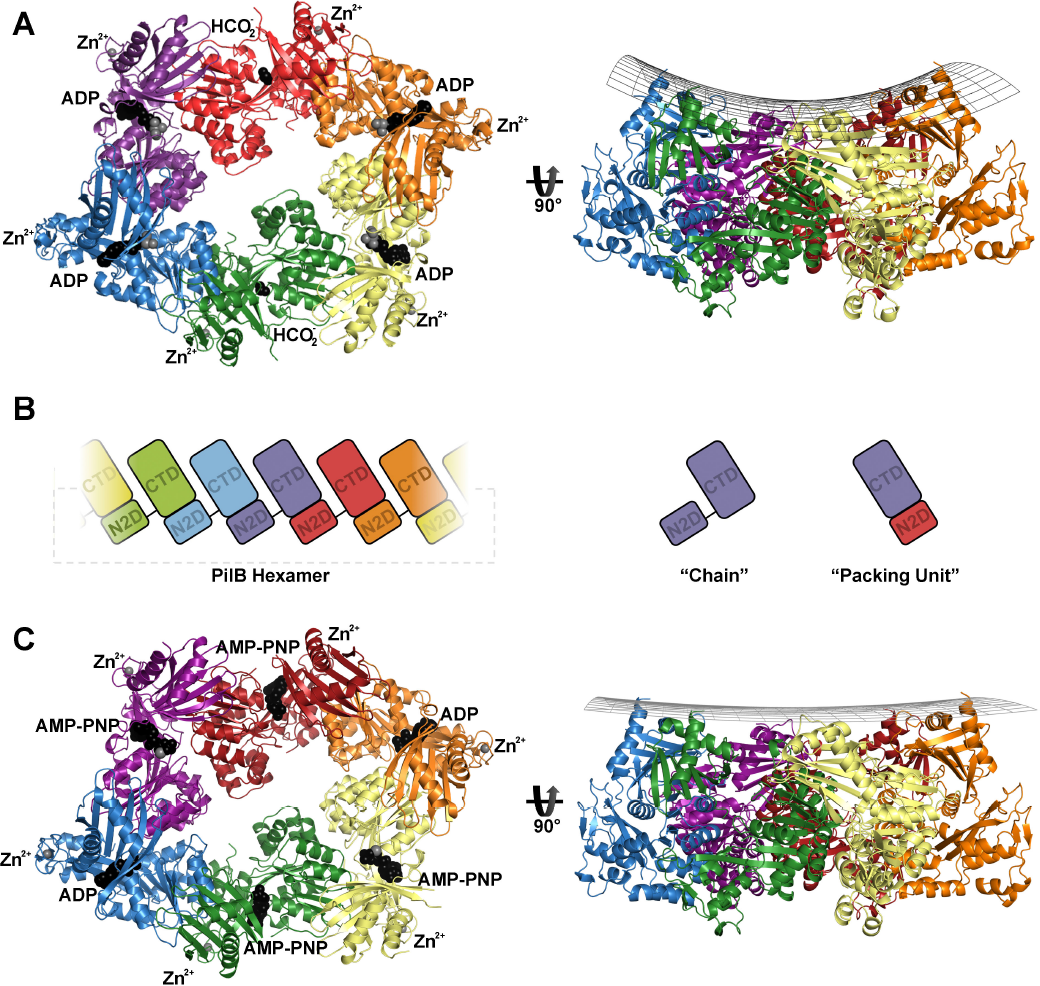
Structure of PilB. (A) PilB:ADP hexamer, with each chain a different color. ADP and formate are shown as black spheres, while magnesium and zinc are shown as grey spheres. A side view is shown with a grid drawn to emphasize the saddle-like shape. (B) Cartoon block illustrations of the PilB hexamer demonstrating the packing units observed, as well as defining the terminology used herein. (C) PilB:AMP-PNP hexamer, with each chain colored as in panel (A). AMP-PNP and ADP are shown as black spheres, while magnesium and zinc are shown as grey spheres. A side view is shown with a grid drawn to emphasize the planar shape.

The principle inter-chain contact, with a buried surface area of ~1600Å, results from interaction between the N2D and the CTD of adjacent chains. This interaction links the N2D of one chain to the CTD of an adjacent chain as a single packing unit, as was noted for GspE, PilT, and FlaI [12, 17-20, 24]. A flexible linker between the N2D and CTD covalently connects two packing units, creating a hexamer of packing units, similar to six beads on a string (FIGURE 2B). Henceforth, where relevant, we refer to these packing units instead of individual chains.

### Structure determination of PilB bound to non-hydrolysable ATP analogue

To trap PilB in a conformation similar to the ATP-bound state, PilB was incubated with AMP-PNP, a non-hydrolysable ATP analogue, under conditions similar to those that facilitated crystallization of PilB bound to four ADP molecules. Crystals formed when AMP-PNP was added to a final concentration of 50 µM, and diffracted to 2.3Å in all three dimensions. The structure was solved by molecular replacement, and again—despite starting with full-length PilB—the N1D of PilB could not be built into the electron density. Six chains could be built in the asymmetric unit, which together formed an elongated hexamer similar to the previously identified PilB hexamer (FIGURE 2C). Henceforth, the AMP-PNP and ADP bound hexamers are referred to as PilB:AMP-PNP and PilB:ADP, respectively.

Like the PilB:ADP hexamer, there was density in the predicted ATP binding sites of all six packing units. However, there were significant differences in the PilB:AMP-PNP hexamer. In two packing units, AMP-PNP, magnesium, and several highly ordered waters best fit the electron density. In two other packing units, ADP with ~50% occupancy best fit the electron density. In the final two packing units, AMP-PNP with ~70% occupancy best fit the electron density.

### The pore of the PilB hexamer is a conserved and negatively charged

Assessing the phylogenetic conservation and surface electrostatics of PilB revealed that there is a conserved and negative surface in the pore of the PilB hexamer (FIGURE 3). The elongated shape of the structure gives the pore the appearance of two ~24 Å diameter sub-pores. The perimeter of the pore is less conserved and positively charged. It is expected that PilC binds the pore of PilB [15], so we extended the aforementioned phylogenetic analysis to residues in the pore of PilB to reveal patterns of conservation between different PilT-like ATPases. Residues in pore loops 2 and 3 are conserved within sub-families but not between sub-families, while D398 or E400 of pore loop 1 is conserved across most PilT-like ATPases (FIGURE 1B). Of note, there are six and seven additional residues in pore loop 3 of the PilT/PilU clade and BfpF clade, respectively.

**Figure 3.**
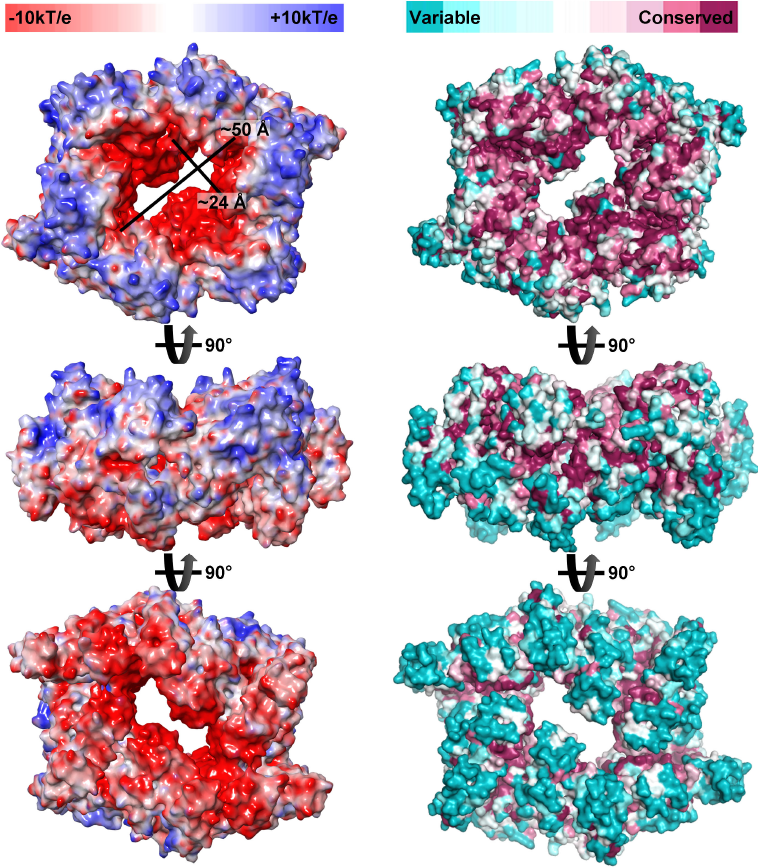
Surface representations of PilB from *G. metalloreducens*. (Left) The electrostatic surface of PilB calculated with missing side chains added in the most likely rotamer via Maestro (version 10.5, Schrödinger). (Right) The phylogenetic conservation of PilB residues mapped onto the surface of PilB using the ConSurf server [51].

### Nucleotide binding correlates with large conformational differences in PilB

In PilB:ADP, ADP is bound between two packing units. In one packing unit, packing unit 1 for simplicity, binds ADP via the Walker A motif and magnesium via E357 coordination, while the phosphates of ADP and E357 bind the adjacent packing unit, packing unit 2, via R286 and R271, respectively (FIGURE 4A and 4B). E357 of the ASP box assumes the role of magnesium coordination typically played by the Walker B motif in ASCE ATPases. At this interface, the backbone of T327 from packing unit 1 contacts the backbone of T411 in packing unit 2. T411 is sandwiched between H420 and H413 of the HIS box.

**Figure 4.**
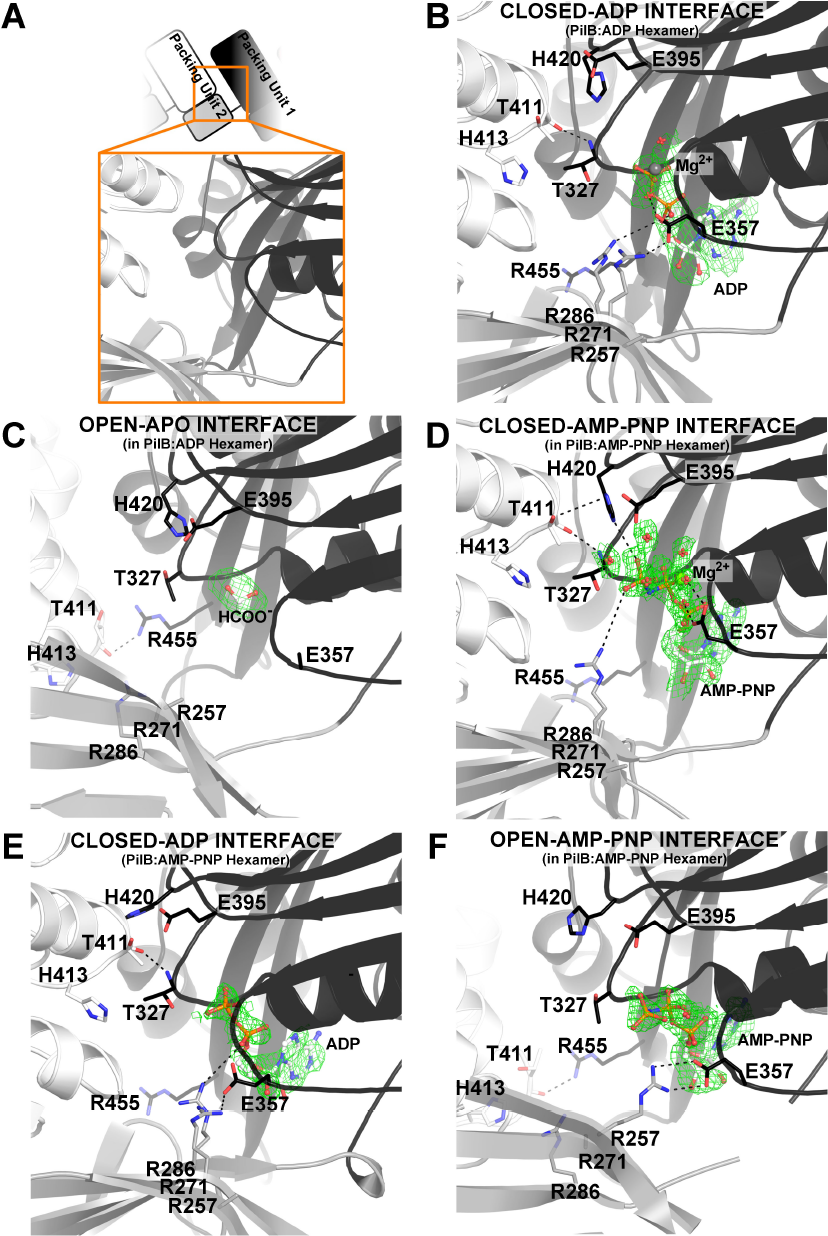
ATP binding sites of PilB. Direct polar contacts are shown as dashed lines. Magnesium is shown as a grey sphere. The green mesh represents the Feature Enhanced Map computed by PHENIX-FEM[52] contoured at 2.0σ. (A) Cartoon clarifying the identity of domains in the following sub-figures. The cartoon mirrors Figure 2C. (B) Nucleotide binding site in the closed-ADP interface from PilB:ADP. (C) Nucleotide binding site in the open-APO interface from PilB:ADP. (D) Nucleotide binding site in the closed-AMP-PNP interface from PilB:AMP-PNP. (E) Nucleotide binding site in the closed-ADP interface from PilB:AMP-PNP. (F) Nucleotide binding site in the open-AMP-PNP interface from PilB:AMP-PNP.

In PilB:ADP there is also an interface between two packing units that is not mediated by ADP. At this interface the orientation of the packing units does not facilitate the binding of T327 to T411; instead, the side chain of R455 of packing unit 1 binds the backbone of T411 of packing unit 2, while the flexible linker plays a more active role in tethering packing units together (FIGURE 4C). Comparing this apo interface to the ADP-bound interface suggests that two packing units close by ~60° to pinch the nucleotide [25] (Supplemental 2). Henceforth, these interfaces will be referred to as the closed-ADP and open-APO interfaces.

The closed-ADP interface has helical character (Supplementary 3), and thus six packing units connected by such interfaces would not create a hexamer but rather a helix with approximately −12 Å rise and 65° twist. Likewise, the open-APO interface also has helical characteristics, but with approximately +24 Å rise and 76° twist. With four closed-ADP interfaces, and two open-APO interfaces, the net rise is zero; and thus six packing units form a ring instead of a helix. This scenario implies conformational restraints on the hexamer to maintain a closed ring: every two closed-ADP interfaces require an open interface to counter the change in rise. This pattern of one open interface for every two closed-ADP interfaces gives the hexamer its elongated appearance. The net twist applied by these interfaces is greater than 360°, so in addition to being elongated, the hexameric ring puckers, similar to the boat conformation of cyclohexanes (FIGURE 2A).

### AMP-PNP binding induces a distinct closed interface

While both PilB:ADP and PilB:AMP-PNP have four closed and two open interfaces, PilB:AMP-PNP is distinct in that only two closed interfaces have ADP bound. The other two closed interfaces are bound to AMP-PNP, henceforth referred to as the closed-AMP-PNP interface. AMP-PNP is also bound in the two open interfaces and will be referring herein as the open-AMP-PNP interface.

The polar contacts created by the closed-ADP and open-AMP-PNP interfaces in the PilB:AMP-PNP hexamer were very similar to the closed-ADP and open-APO interfaces in the PilB:ADP hexamer, respectively (FIGURE 4E and 4F). However, in the open-AMP-PNP interface, E357 of packing unit 1 forms a salt bridge with R257 of packing unit 2. This interaction did not occur in the open-APO interface of PilB:ADP.

The closed-AMP-PNP interface is also similar to that of the closed-ADP interface of PilB:ADP. The backbone of T327 from packing unit 1 still contacts the backbone of T411 from packing unit 2. However, H420 of the HIS box coordinates the ɣ-phosphate of AMP-PNP and the side chain of T411 from packing unit 2, suggesting that the HIS box plays a role in mediating the inter-chain cooperativity of ATP catalysis. In addition, since the magnesium at the closed-AMPPNP interface contacts the β- and ɣ-phosphate, E357 is reoriented to coordinate magnesium. In this position, E357 from packing unit 1 cannot bind R271 from packing unit 2. Nearby, R286 coordinates the ɣ-phosphate of the AMP-PNP instead of coordinating the α-phosphate of ADP (FIGURE 4D). In this way, R286 and R271 in packing unit 2 sense the presence of the ɣ-phosphate bound to packing unit 1, and twist the closed-AMP-PNP interface by 8° relative to the closed-ADP interface [25].

The helical rise and twist created by the closed-AMP-PNP interface is approximately −12 Å and 62°, respectively, which is accommodated by the open interface that applies approximately +24 Å rise and 72° twist. The closed-ADP interface still has approximately −12 Å rise and 65° twist. The net decrease in twist reduces the ring puckering relative to the boat shape of PilB:ADP such that the PilB:AMP-PNP ring appears more planar (FIGURE 2C).

### Heterogeneous nucleotide binding indicates catalytic mechanism

Even through no exogenous nucleotide was added during crystallization, PilB:ADP crystallized with four ADP molecules. These nucleotides must have been pull-down from the cytoplasm of *E. coli* during purification. It is possible that ATP was also pulled down but it was subsequently catalyzed to ADP. The absence of nucleotide in the open interface suggests that this site has a relatively low affinity for ADP compared with the closed interfaces. Also, the presence of ADP at the closed-ADP interfaces indicates that PilB does not immediately release ADP following ATP catalysis. ADP release from the closed-ADP interface may be contingent on a separate event, such as ATP binding to the open-APO interface.

During crystallization of PilB:AMP-PNP AMP-PNP was limited with 50 µM AMP-PNP added to 120 µM solution of PilB, a ratio of 5:12 nucleotides to ATP binding sites. As a result, the occupancy of AMP-PNP was not uniform across the PilB hexamer. The two closed-AMPPNP interfaces bound AMP-PNP with full occupancy, the two open-AMP-PNP interfaces bound AMP-PNP with partial occupancy, while the two closed-ADP interfaces bound ADP with partially occupancy not AMP-PNP. This suggests that the two closed-AMP-PNP interfaces have the highest affinity for AMP-PNP, and by extension ATP, while the open interfaces have a moderate affinity for ATP, and the closed-ADP interfaces have low affinity for ATP. The partial occupancy of ADP at the closed-ADP interface of PilB:AMP-PNP is consistent limited nucleotide binding to the open interface triggering ADP release from the closed-ADP interface.

Together the PilB:ADP and PilB:AMP-PNP structures indicate the relative affinities of ATP and ADP for the different interfaces observed in the PilB hexamer. The open interface has a low affinity for ADP and a moderate affinity for ATP. With the N2Ds facing the viewer, the closed interface immediately clockwise of the open interface has a high affinity for ADP and a low affinity for ATP. The next closed interface has a high affinity for ATP and ADP. This provides evidence for the mechanism of ATP turnover. Starting with the PilB:ADP hexamer— PilB bound to four ADP molecules—added ATP would bind the two open interfaces since the closed interfaces are occupied with ADP. ATP binding to the open interface would result in the closure of this interface, and because of the aforementioned rise restraints on the hexamer, this would trigger opening of two closed-ADP interfaces and the release ADP. The location of the closed-AMP-PNP and closed-ADP interfaces in the PilB:AMP-PNP hexamer demonstrates that there is directionality to the release of ADP. ADP is released from the closed interfaces immediately clockwise from the open interfaces that bind ATP. Finally, PilB would be reset after ATP catalysis yielding PilB bound to four ADP molecules.

### Modeling ATP turnover in PilB suggests pore movements

To model the structural changes in PilB that may occur as ATP is catalyzed to ADP, we created an interpolated trajectory of the PilB:AMP-PNP structure to the PilB:ADP structure. In this model, one can visualize that ATP catalysis may cause the relatively planar PilB hexamer to adopt a saddle shape (ANIMATION 1). We also modeled the structural change in PilB that may occur as a new ATP is bound and ADP is released; to do this we created an interpolated trajectory of the PilB:ADP structure to the PilB:AMP-PNP structure (ANIMATION 2). The model reflects the expectation that ATP binding induces closure of two open interfaces, while opening two of the closed interfaces. In this model, ATP binding causes two motions perpendicular to one another in the PilB hexamer: a clockwise rotation of the sub-pores and a rotation of the two packing units that went from being closed to open (FIGURE 5A and ANIMATION 3). With the N2Ds facing the viewer, the sub-pore rotation is clockwise and the packing units scoop towards the viewer. The conserved pore residue E400 on the scooping packing units moves towards the viewer by ~13 Å.

**Figure 5.**
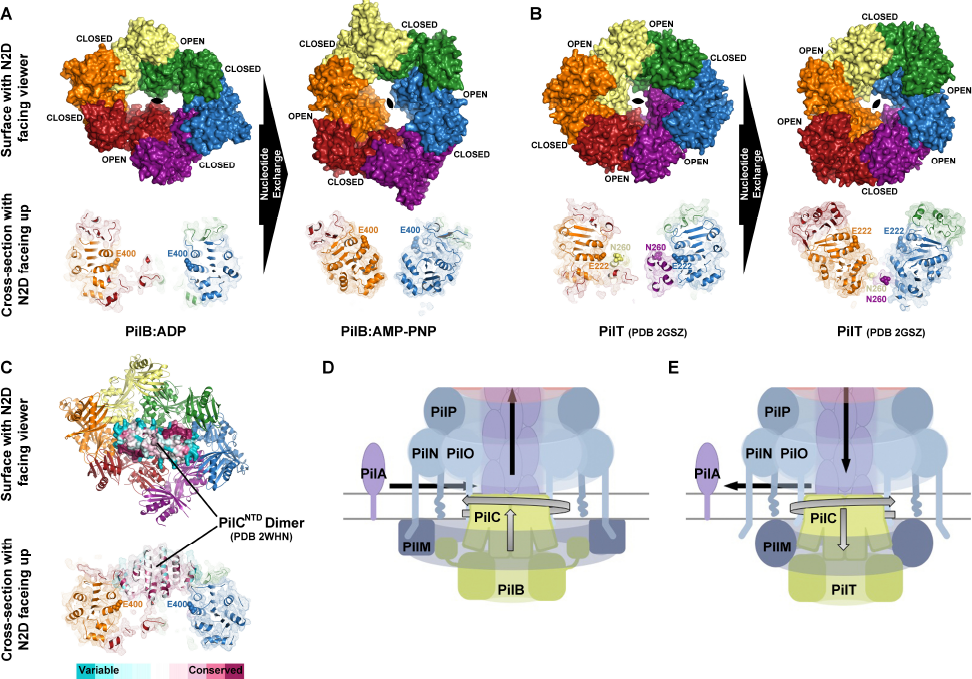
Modeling the movements and functions of PilB and PilT ATPases. **(A)** The PilB:ADP hexamer was aligning with the PilB:AMP-PNP hexamer to suggest the movements that may occur as two closed packing units open and two open packing units close during nucleotide exchange. Promoters are colored as in Figure 2. (Bottom) A cross-section through the center of PilB is shown with E400 displayed as spheres. **(B)** The PilT hexamer (PDB 2GSZ) was aligned with the same PilT hexamer rotated by one packing unit to suggest the movements that may occur as two closed packing units open and two open packing units close during nucleotide exchange. (Bottom) A cross-section through the center of PilT is shown with E222 (the equivalent of E400 in PilB) and N260 from the midpoint of the extension on pore loop 3 (there is no equivalent in PilB) displayed as spheres. **(C)** The PilC^NTD^ dimer (PDB 2WHN) was manually placed in the two sub-pores of PilB, and the phylogenetic conservation of PilC residues were mapped onto the surface using the ConSurf server [51]. (Bottom) A cross-section through the center of PilB is shown with E400 displayed as spheres. The model is similar to the mode of PilC binding to PilB we proposed previously [15]. **(D)** Working model for the molecular mechanism of the PilB motor. We propose that PilC is pushed by PilB upwards towards the membrane, allowed to fall back, and rotated in 60° increments to facilitate helical PilA polymerization. **(E)** Working model for the molecular mechanism of the PilT motor. We propose that PilC is pulled by PilB downward towards the cytoplasm, allowed to fall back, and rotated in 60° increments in the opposite direction of PilT to facilitate helical PilA depolymerization.

### The same direction of ATP turnover in PilT produces opposite pore movements relative to PilB

We next compared the PilB structures to the C2 symmetric PilT hexamer, published previously [17]. By structural comparison with the interfaces of PilB, we identified two closed and four open interfaces in PilT (Supplemental 3). PilB and PilT appear to have an enantiomeric arrangement of open and closed interfaces (Figure 5B).

Based on the direction of ATP turnover in PilB, we modelled the structural changes that may occur as ATP is turned over in PilT. Similar to the animated PilB model, the animated PilT model reflects the expectation that nucleotide exchange induces closure of two open interfaces and opening two of the closed interfaces. Like PilB, this causes two motions perpendicular to one another in the PilT hexamer: a rotation of the pores, and a rotation of the two packing units that went from being closed to open (Figure 5B and ANIMATION 4). Despite assuming the same direction of ATP turnover as PilB, with the N2D facing the viewer the rotation of the PilT pore is counter clockwise. Similar to PilB, in PilT there is a rotation of the two packing units that went from being closed to open. However, in PilT these packing units have limited access to the pore due to steric hindrance from the extension of residues on pore loop 3 from another packing unit in the hexamer. Notably, the residues in this extension sweep away from the viewer by ~19Å. In other words, if we assume the same direction of ATP turnover in PilB and PilT, opposite pore movements result as a consequence of two unique adaptions in PilT: the enantiomeric arrangement of open and closed interfaces, and an extension of residues on pore loop 3.

## DISCUSSION

While this paper was being completed, a C2 symmetric structure of *Thermus thermophilus* PilB (PilB_Tt_) at 2.7 Å resolution was published [26]. Despite the coordinates of this structure not being currently available, comparisons can be made. In contrast to our structures of PilB from *G. metalloreducens* (PilB_Gm_), the PilB_Tt_ hexamer is homogeneously saturated with a non-hydrolysable ATP analogue, similar to the structures of FlaI and archaeal GspE2. In PilB_Tt_ a solvent channel was identified leading to an interface equivalent to the closed-AMP-PNP interface of the PilB_Gm_:AMP-PNP structure presented herein [26]. This channel was proposed to facilitated ADP/ATP exchange [26]. In addition, in the closed interfaces of PilBTt, the ɣ-phosphate of the ATP analogue was modeled into the position occupied by magnesium in the open interface of PilB_Tt_. Likewise, magnesium could not be modelled into the closed interfaces of PilB_Tt_, and it was proposed that the closed interfaces of PilB_Tt_ would not hydrolyse ATP. [26]. Thus based on PilB_Tt_, the direction of ATP turnover proposed is the opposite to the direction proposed herein. Likewise, the PilB_Tt_ pore was suggested to rotate counter clockwise while a different pair of packing units scooped through the pore.

The increased resolution and the heterogeneous nucleotide binding observed in our PilB_Gm_ structures enables improved annotation of the nucleotide binding sites and revealed a correlation between nucleotide binding and inter-chain interfaces. PilB_Gm_ is the only PilT-like ATPase bound to an ATP analogue and magnesium that has been crystallized at a resolution sufficient to differentiate the octahedral water coordination of magnesium from the tetrahedral ɣ-phosphate. This has enabled us to define with more confidence the direction of ATP turnover and more thoroughly illustrate the conformational changes that occur. The closed-AMP-PNP interface of the PilB_Gm_:AMP-PNP structure is the only interface with the catalytic glutamate poised for hydrolysis indicating that this is the site of ATP catalysis, not nucleotide exchange. In addition, the vacancy and partial occupancy of nucleotide in the open interfaces of the PilB_Gm_:ADP structure and PilB_Gm_:AMP-PNP structure, respectively, indicate that this site is used for ADP/ATP exchange not for ATP catalysis. With this knowledge, we built a model for PilB movements distinct from the PilB_Tt_ model, and then expanded our analysis to the related ATPase, PilT.

We also identified pore residues that are highly conserved in PilT-like ATPases. As previously hypothesized, PilC^NTD^ is expected to bind in the pore of PilB [15], and thus these residues may be important for interaction with PilC. A crystal structure of PilC^NTD^ from *T. thermophilus* was solved as an asymmetric dimer [27]. The diameter of each PilC^NTD^ chain is approximately 23Å, and thus the PilC^NTD^ dimer would fit snugly in the two ~24Å sub-pores of PilB (FIGURE 5C). Examining the phylogenetic conservation of the PilC^NTD^ dimer in the context of our model of PilB hexamer movements revealed a patch of conserved residues in PilC^NTD^ adjacent to the packing units that we predict scoop laterally through the pore. With this in mind, we propose that PilB functions by ratcheting the PilC^NTD^ dimer upwards into the inner membrane. Following the rotary motions of the PilB sub-pores as ATP is hydrolyzed, PilC may turn clockwise at 60° increments. By the end of the 60° rotation, the PilC^NTD^ dimer will no longer be oriented to bind the packing units that swept through the pore. Thus, we propose that PilC is pushed upwards towards the membrane, allowed to fall back, and rotated by 60° in a single motion, like a revolver, (FIGURE 5D). There are several lines of evidence indicating that PilC binds directly to PilA [16, 28, 29], and thus this ratcheting action could facilitate PilA extraction from the membrane and insertion into the pilus. If a pilin were inserted with every ratcheting event, the pilus would be a right-handed helix with a 60° twist. This is consistent with reports in the literature that suggest the pilus is a right-handed helix with a range of twists (60-100°) within and between systems and species [30-34], and thus we predict that the pilus itself facilitates further twisting upon assembly—as predicted by molecular dynamic simulations [30]. Of note, the model of PilB movements described based on the PilB_Tt_ structure would build a left-handed helix, and the patch of conserved residues on PilC^NTD^ would not align with the packing units suggested to scoop through the pore.

Applying similar transformations to the available structure of C2 symmetric PilT suggested the structural changes that may occur as nucleotide is exchanged. Despite applying the same direction of opening and closing interfaces, the rotation of the PilT pore would be opposite that of PilB. This is the direct consequence of the enantiomeric arrangement of open and closed interfaces in PilB and PilT (Supplemental Figure 4). Furthermore, a conserved extension on pore loop 3 sweeps laterally through the pore in the opposite direction of the scooping motion in PilB. We predict that PilC binds to the extension on pore loop 3, and propose that PilT acts an anti-PilB: rotating PilC in the opposite direction of PilB and pulling PilC towards the cytoplasm to facilitate PilA depolymerisation (FIGURE 5E).

While the proposed PilB and PilT mechanisms rely on C2 symmetry with restricted open and closed interfaces, we cannot omit the possibility of alternative interfaces not characterized here, as there are models of FlaI and archaeal GspE2 with C3 symmetry. That FlaI and archaeal GspE2 would form similar hexamers is not surprising given their clustering in our phylogenetic analysis. It is interesting to note however that unlike other PilT-like ATPases, FlaI could be responsible for two rotary functions: turning FlaJ to power assembly as outlined for PilB [35], and then spinning the assembled archaeal flagellum. If the C2 rotary model is responsible for the former, it is tempting to speculate that the unique C3 symmetry of FlaI could reflect a functional state involved in turning the archaeal flagellum once assembly is complete.

An assumption implicit in the C2 rotary model is the orientation of PilB in relation to the inner membrane and the T4aP. We based this assumption on the electrostatic surface properties of PilB: the positive face on the PilB hexamers may correspond to an interface used to bind the inner membrane. Indeed, electron tomography studies that shows PilB adjacent to the cytoplasmic face of the inner membrane [16], and the PilB-like ATPase, GspE, has increased ATPase activity in the presence of cardiolipins, suggesting that it binds directly to the inner membrane [36].

We also noted that PilB:ADP adopted a saddle shape, while the AMP-PNP bound conformation was relatively planar. This architecture implies that upon ATP catalysis, the inner membrane may be deformed, and/or that the inner membrane applies a strain on these saddle-shaped hexamers. This strain would then be released as the hexamer bound more ATP and ratcheted PilC, adopting a planar state. Decoupling the energy-generating step from the energy-using step using strain is a common strategy in macroscopic biology to amplify force [37, 38]. Thus, it is possible that PilB temporarily stores strain to ratchet PilC with amplified force. If PilT acts likewise, it would help explain why the pilus can retract with the extraordinary forces for which it is renowned.

## ACKNOWLEDGEMENTS

This work was supported by a grant from the Canadian Institutes for Health Research (CIHR) MOP 93585 to LLB and PLH. MM is supported by a CIHR doctoral studentship. PLH is the recipient of a Canada Research Chair. Funds for the X-ray facilities at The Hospital for Sick Children were provided, in part, by the Canadian Foundation for Innovation.

## EXPERIMENTAL PROCEDURES

### Cloning, Expression, and Purification of PilM, PilB, and PilT

PilB from *G. metalloreducens* was PCR amplified from genomic DNA using primers P96 (TATATATAGCTAGCATGCAGGCCAGCAGACTG) and P97 (TATATATAGGATCCTTAATCGTCGGCCACGGTG), digested with NheI and BamHI, and cloned into pET28a with a thrombin-cleavable hexahistadine tag to create pET28a:PilB^Gm^. The fidelity of the sequence was verified by TCAG sequencing facilities (SickKids, Canada). *Escherichia coli* BL21-CodonPlus® cells (F^−^ *ompT hsdS* (rB^−^ mB^−^) dcm^+^ Tet^r^ galλ (DE3) endA [*argU proL* Cam^r^]; Stratagene, USA) transformed with pET28a:PilB^Gm^, grown in 4 L of lysogeny broth (LB) with 100 µg/ml kanamycin at 37 °C to an A600 of 0.5–0.6. Protein expression was induced by the addition of isopropyl-D-1-thiogalactopyranoside (IPTG) to a final concentration of 1 mM, and the cells were grown for 16 h at 18 °C. Cells were pelleted by centrifugation at 9000 *× g* for 15 min. Cell pellets were subsequently resuspended in 40 ml binding buffer (50 mM Tris pH 7.5, 150 mM NaCl, and 50 mM imidazole), lysed by passage through an Emulsiflex-c3 high-pressure homogenizer, and the cell debris removed by centrifugation for 60 min at 40000 *× g*. The resulting supernatant was passed over a column containing 5 ml of pre-equilibrated Ni-NTA agarose resin (Life Technologies, USA). The resin was washed with 10 column volumes (CV) of binding buffer and eluted with binding buffer plus 300 mM imidazole. The protein was then further purified by size exclusion chromatography on a HiLoadTM 16/600 SuperdexTM 200pg column pre-equilibrated with binding buffer without imidazole. Purified proteins were stored at 4°C for less than 2 days before use.

### Crystallization, Data Collection, and Structure Solution

For crystallization, purified PilB was concentrated to 16 mg/ml at 3000 *× g* in an ultrafiltration device (Millipore). Crystallization conditions were screened using the complete MCSG suite (MCSG 1-4) (Microlytic, USA) using a Gryphon LCP robot (Art Robbins Instruments, USA). Crystal conditions were screened and optimized using vapour diffusion at 20 °C with Art Robbins Instruments Intelli-Plates 96-2 Shallow Well (Hampton Research, USA) with 1 µl protein and 1 µl reservoir solution. For the PilB:AMP-PNP structure, the protein solution also contained 100 µM AMP-PNP. For the PilB:ADP structure the reservoir solution was 13% (w/v) PEG3350, 0.1 M magnesium formate, 0.1 M Tris pH 7.6. For cryoprotection, 2 µl of 1:1 ethylene glycol and reservoir solution was added to the drop containing the crystal for 10 seconds prior to vitrification in liquid nitrogen. For the PilB:AMP-PNP structure, the reservoir solution was 11% (w/v) PEG3350, 0.1 M magnesium formate, 0.1 M Tris pH 7.0. For cryoprotection, 2 µl of 1:1 2-methyl-2,4-pentanediol and reservoir solution was added to the drop containing the crystal for 10 seconds prior to vitrification in liquid nitrogen. Diffraction data was collected on Beamline 08ID-1 at the Canadian Macromolecular Crystallography Facility (see Table 1). The PilB:AMP-PNP data were indexed, scaled, and truncated to 2.3 Å using XDS [39]. The PilB:ADP data were indexed in XDS, then anisotropically truncated to 3.4 Å in one dimension and 3.9 Å in the other two dimensions and scaled using the Diffraction Anisotropy Server [40]. PHENIX-MR [41] was used solve the structure of PilB by molecular replacement with residues 100-226 and residues 236-500 of GspE (PDB 1P9R) pre-processed by the program Chainsaw [42]. The resulting electron density maps were of high quality and enabled building the PilB proteins manually in COOT [43]. Through iterative rounds of building/remodelling in COOT [43] and refinement in PHENIX-refine [44] the structures of the PilB:ADP and PilB:AMP-PNP were built and refined. PilB:ADP was refined with group B-factors, while PilB:AMP-PNP was refined with individual B-factors. The occupancy of the nucleotides in the PilB:AMP-PNP structure was refined such that all atoms in a nucleotide had the same occupancy. Progress of the refinement in all cases was monitored using Rfree.

**Table 1.** Data Collection and Refinement Statistics of PilM and PilM:PilN_1-12_ Structures. Note: Values in parentheses correspond to the highest resolution shell.

|  | PilB:ADP | PilB:AMP-PNP |
| --- | --- | --- |
| **Data Collection** |  |  |
| Beamline | 08ID-1 | 08ID-1 |
| Wavelength (Å) | 0.97920 | 0.97920 |
| Space group | *P*3<sub>2</sub>21 | *P*1 |
| *a*, *b*, *c* (Å) | 112.1, 112.1, 299.2 | 74.3, 100.0, 109.8 |
| $\alpha$, $\beta$, $\gamma$ (°) | 90.0, 90.0, 120.0 | 113.0, 107.7, 88.9 |
| Resolution (Å) | 46-3.40<br>(3.52-3.40) | 48-2.30<br>(2.38-2.30) |
| Total Reflections | 208303 | 253982 |
| Unique Reflections | 23642 (375) | 120979 (1531) |
| Redundancy | 8.8 (10.3) | 2.1 (2.1) |
| Completeness (%) | 77 (12)* | 96 (95) |
| Mean *I* / $\sigma I$ | 12.7 (2.78) | 5.8 (2.0) |
| *R*<sub>Sym</sub> (%)** | 21 (94) | 5.6 (43) |
| CC* (%)** | 100 (96) | 100 (88) |
| **Refinement** |  |  |
| *R*<sub>work</sub> / *R*<sub>free</sub> (%)† | 27.2 / 28.9 | 18.1 / 22.0 |
| RMSD |  |  |
| Bond lengths (Å) | 0.007 | 0.011 |
| Bond angles (°) | 1.42 | 0.91 |
| Ramachandran plot‡ |  |  |
| Total favoured (%) | 98 | 99 |
| Total allowed (%) | 100 | 100 |
| Coordinate error (Å)§ | 0.38 | 0.27 |
| Atoms | 8477 | 18174 |
| Protein | 8406 | 17443 |
| Water | 12 | 533 |
| Magnesium | 2 | 6 |
| AMP-PNP | 0 | 132 |
| ADP | 54 | 54 |
| Zinc | 3 | 6 |
| Av. B-factors (Å<sup>2</sup>)‡ | 87.7 | 49.3 |
| Protein | 87.9 | 49.2 |
| Water | 36.7 | 48.2 |
| Ligands | 65.6 | 63.2 |

\* atypical completeness of the PilB:ADP structure reflects anisotropic truncation
\*\* $R_{\text{Sym}} = \sum \sum |I - \langle I \rangle| / \sum \sum I$ , $R_{\text{pim}} = \sum \sqrt{(1/(n-1)) \sum |I - \langle I \rangle| / \sum \sum I}$ , and $\text{CC}^* = \sqrt{(2\text{CC}_{1/2}/(1+\text{CC}_{1/2}))}$ where $\text{CC}_{1/2}$ is the Pearson correlation coefficient of two half data sets as described elsewhere [49]
† $R_{\text{work}} = \sum ||F_{\text{obs}}| - k|F_{\text{calc}}|| / |F_{\text{obs}}|$ where $F_{\text{obs}}$ and $F_{\text{calc}}$ are the observed and calculated structure factors, respectively. $R_{\text{free}}$ is the sum extended over a subset of reflections (5%) excluded from all stages of the refinement
‡ As calculated using MolProbity [50]
§ Maximum-likelihood based Coordinate Error, as determined by PHENIX [41]

### Phylogenetic analysis

HMMER [45] was used to identify PilB orthologues. The redundancy of the orthologues was reduced to <40% while retaining the orthologues from model systems. The remaining sequences were aligned in ClustalOmega [46], and a Neighbour-Joining tree was built with MEGA using the Poisson substitution model [47]. Sequence logos were generated using WebLogo [48].

---

**Figure S1.**
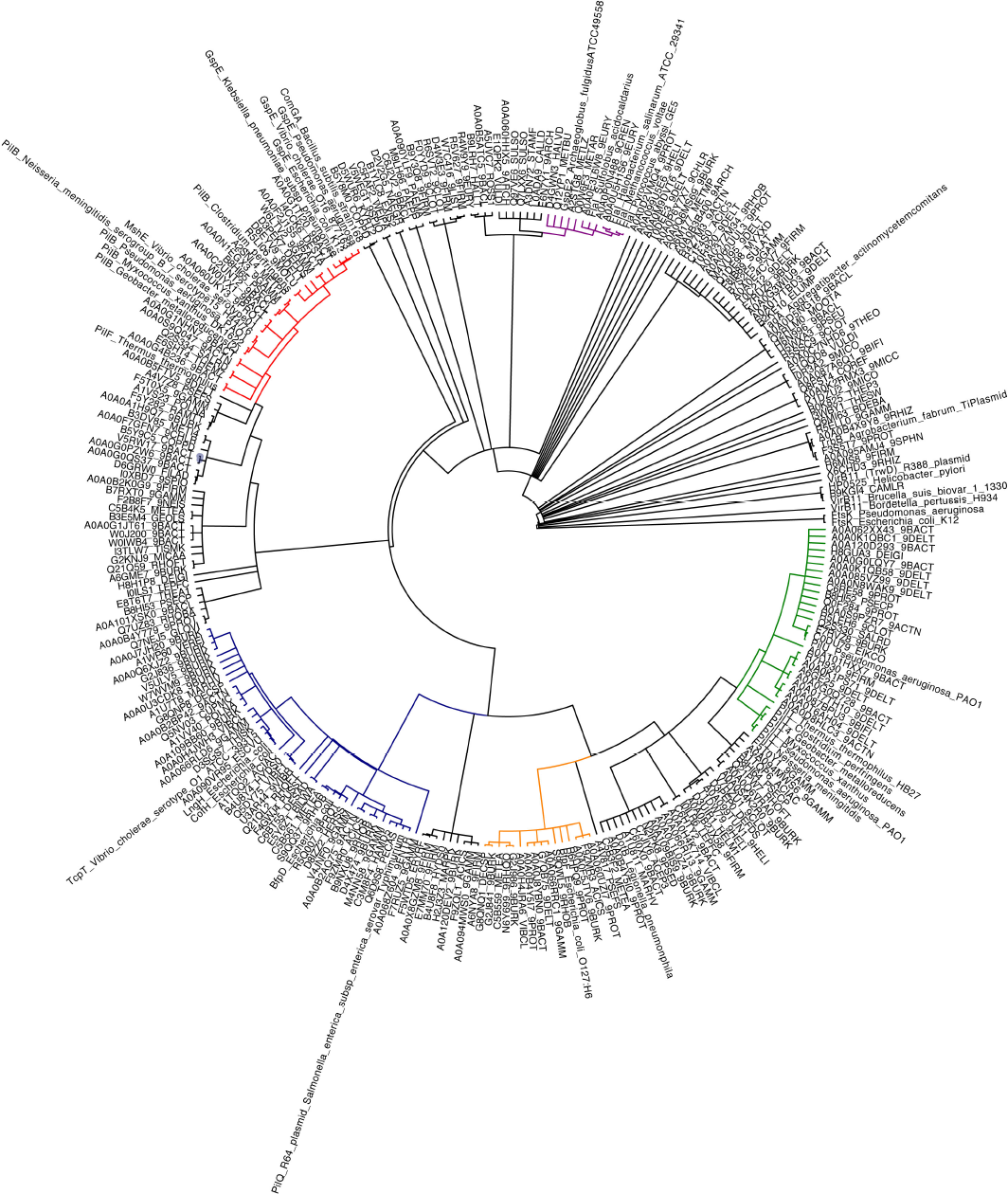
Phylogenetic analysis of PilT-like ATPases. Sequences are labeled with their PFAM identifiers, or with the protein name and species if the sequence is from a model system. Only branches with a >85% bootstrap value are shown (1000 bootstraps). FtsK was used as an out-group. Protein sub-families, or clades, are given a unique color for clarity and to stratify the sequences for further analysis. Red is for the PilB/GspE clade. Purple is for the FlaI/GspE2 clade. Green is for the PilT/PilU clade. Orange is for the BfpF clade. Blue is for the BfpD/TcpT clade.

**Figure S2.**
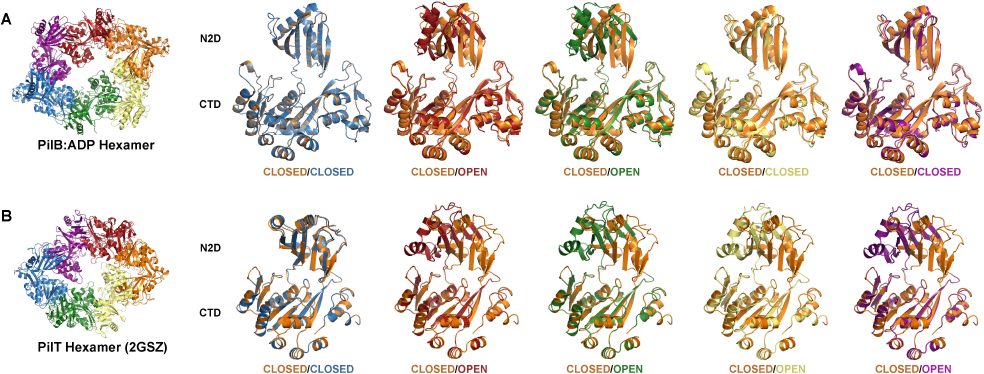
Open or closed arrangement of interfaces in PilB and PilT. Hexamers are colored as in Figure 2. Individual chains are aligned with the C2D of a closed chain (orange) for comparison. Since the N2D and CTD of a chain belong to adjacent packing units, individual chains are the most convenient way to display angle changes between interfaces.

**Figure S3.**
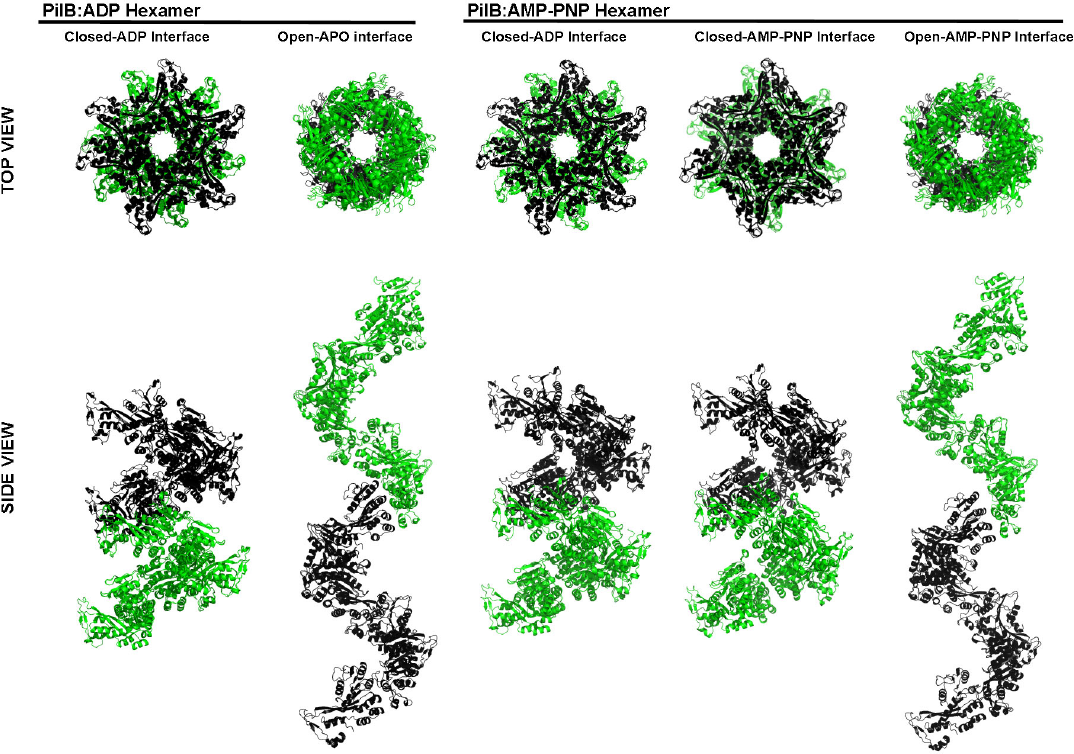
Helical properties of the packing unit interfaces in PilB. Twelve packing units were aligned to simulate a helix formed by the specified PilB interfaces. For simplicity, only the C2D of each packing unit is shown, the first six units are coloured black, while the last six are coloured green. The twist and rise of the closed-ADP interface from the PilB:ADP hexamer and the PilB:AMP-PNP hexamer was ~65° and -12 Å, respectively. The twist and rise of the open-APO interface from the PilB:ADP hexamer was ~76° and +24 Å, respectively. The twist and rise of the closed-AMP-PNP interface from the PilB:AMP-PNP hexamer was ~62° and -12 Å, respectively. The twist and rise of the open-AMP-PNP interface from the PilB:AMP-PNP hexamer was ~72° and +24 Å, respectively.

**Figure S4.**
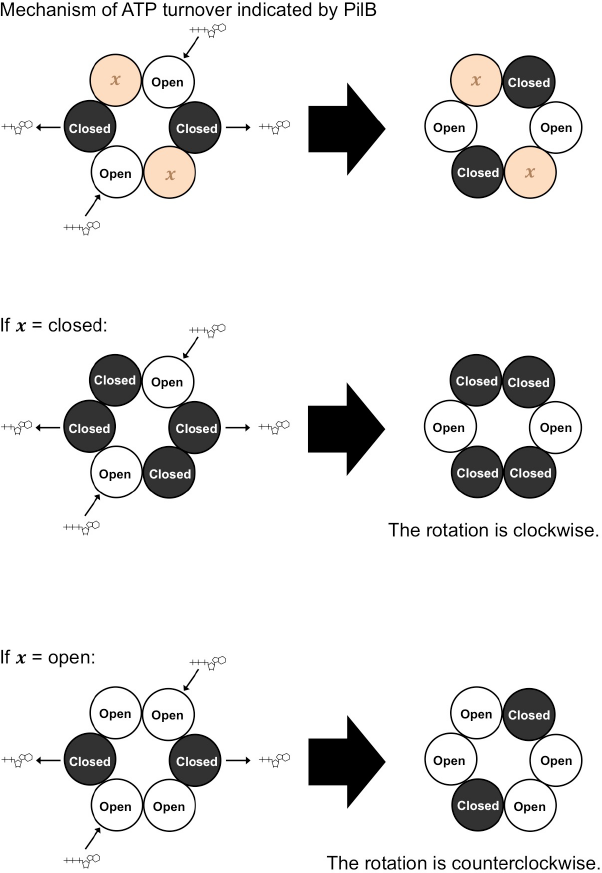
Cartoon describing why the enantimeric arrangement of open and closed interfaces in PilB and PilT leads to opposite directions of elongated pore rotation. ATP and ADP are shown as sticks. The interface between two packing units is shown as a large sphere; if it is closed it is black, if it is open it is white. *Top,* the direction of ATP turnover indicated by the structures of PilB and the resulting changes in interface closure. *Middle*, same as *Top* if four interfaces were closed and two were open, as for PilB. *Bottom*, same as *Top* if four interfaces were open and two were closed, as for PilT.

## REFERENCES

1. Maier B, Potter L, So M, Long CD, Seifert HS, Sheetz MP. Single pilus motor forces exceed 100 pN. Proc Natl Acad Sci U S A. 2002;99(25):16012–7. doi: 10.1073/pnas.242523299. PubMed PMID: 12446837; PubMed Central PMCID: PMCPMC138556.

2. Burrows LL. Pseudomonas aeruginosa twitching motility: type IV pili in action. Annu Rev Microbiol. 2012;66:493–520. doi: 10.1146/annurev-micro-092611-150055. PubMed PMID: 22746331.

3. Peabody CR, Chung YJ, Yen MR, Vidal-Ingigliardi D, Pugsley AP, Saier MH, Jr. Type II protein secretion and its relationship to bacterial type IV pili and archaeal flagella. Microbiology. 2003;149(Pt 11):3051–72. doi: 10.1099/mic.0.26364-0. PubMed PMID: 14600218.

4. Mattick JS. Type IV pili and twitching motility. Annu Rev Microbiol. 2002;56:289–314. doi: 10.1146/annurev.micro.56.012302.160938. PubMed PMID: 12142488.

5. Berry JL, Pelicic V. Exceptionally widespread nanomachines composed of type IV pilins: the prokaryotic Swiss Army knives. FEMS Microbiol Rev. 2015;39(1):1–21. doi: 10.1093/femsre/fuu001. PubMed PMID: 25793961.

6. Planet PJ, Kachlany SC, DeSalle R, Figurski DH. Phylogeny of genes for secretion NTPases: identification of the widespread tadA subfamily and development of a diagnostic key for gene classification. Proc Natl Acad Sci U S A. 2001;98(5):2503–8. doi: 10.1073/pnas.051436598. PubMed PMID: 11226268; PubMed Central PMCID: PMCPMC30167.

7. Iyer LM, Makarova KS, Koonin EV, Aravind L. Comparative genomics of the FtsK-HerA superfamily of pumping ATPases: implications for the origins of chromosome segregation, cell division and viral capsid packaging. Nucleic Acids Res. 2004;32(17):5260–79. doi: 10.1093/nar/gkh828. PubMed PMID: 15466593; PubMed Central PMCID: PMCPMC521647.

8. Walker JE, Saraste M, Runswick MJ, Gay NJ. Distantly related sequences in the alpha- and beta-subunits of ATP synthase, myosin, kinases and other ATP-requiring enzymes and a common nucleotide binding fold. EMBO J. 1982;1(8):945–51. PubMed PMID: 6329717; PubMed Central PMCID: PMCPMC553140.

9. Jakovljevic V, Leonardy S, Hoppert M, Sogaard-Andersen L. PilB and PilT are ATPases acting antagonistically in type IV pilus function in Myxococcus xanthus. J Bacteriol. 2008;190(7):2411–21. doi: 10.1128/JB.01793-07. PubMed PMID: 18223089; PubMed Central PMCID: PMCPMC2293208.

10. Thomsen ND, Berger JM. Structural frameworks for considering microbial protein- and nucleic acid-dependent motor ATPases. Mol Microbiol. 2008;69(5):1071–90. doi: 10.1111/j.1365-2958.2008.06364.x. PubMed PMID: 18647240; PubMed Central PMCID: PMCPMC2538554.

11. Chiang P, Sampaleanu LM, Ayers M, Pahuta M, Howell PL, Burrows LL. Functional role of conserved residues in the characteristic secretion NTPase motifs of the Pseudomonas aeruginosa type IV pilus motor proteins PilB, PilT and PilU. Microbiology. 2008;154(Pt 1):114–26. doi: 10.1099/mic.0.2007/011320-0. PubMed PMID: 18174131.

12. Yamagata A, Tainer JA. Hexameric structures of the archaeal secretion ATPase GspE and implications for a universal secretion mechanism. EMBO J. 2007;26(3):878–90. doi: 10.1038/sj.emboj.7601544. PubMed PMID: 17255937; PubMed Central PMCID: PMCPMC1794398.

13. Merz AJ, So M, Sheetz MP. Pilus retraction powers bacterial twitching motility. Nature. 2000;407(6800):98–102. doi: 10.1038/35024105. PubMed PMID: 10993081.

14. Nunn DN, Lory S. Product of the Pseudomonas aeruginosa gene pilD is a prepilin leader peptidase. Proc Natl Acad Sci U S A. 1991;88(8):3281–5. PubMed PMID: 1901657; PubMed Central PMCID: PMCPMC51430.

15. Takhar HK, Kemp K, Kim M, Howell PL, Burrows LL. The platform protein is essential for type IV pilus biogenesis. J Biol Chem. 2013;288(14):9721–8. doi: 10.1074/jbc.M113.453506. PubMed PMID: 23413032; PubMed Central PMCID: PMC3617274.

16. Chang YW, Rettberg LA, Treuner-Lange A, Iwasa J, Sogaard-Andersen L, Jensen GJ. Architecture of the type IVa pilus machine. Science. 2016;351(6278):aad2001. doi: 10.1126/science.aad2001. PubMed PMID: 26965631.

17. Satyshur KA, Worzalla GA, Meyer LS, Heiniger EK, Aukema KG, Misic AM, et al. Crystal structures of the pilus retraction motor PilT suggest large domain movements and subunit cooperation drive motility. Structure. 2007;15(3):363–76. doi: 10.1016/j.str.2007.01.018. PubMed PMID: 17355871; PubMed Central PMCID: PMC1978094.

18. Misic AM, Satyshur KA, Forest KT. P. aeruginosa PilT structures with and without nucleotide reveal a dynamic type IV pilus retraction motor. J Mol Biol. 2010;400(5):1011–21. doi: 10.1016/j.jmb.2010.05.066. PubMed PMID: 20595000; PubMed Central PMCID: PMCPMC2918248.

19. Lu C, Turley S, Marionni ST, Park YJ, Lee KK, Patrick M, et al. Hexamers of the type II secretion ATPase GspE from Vibrio cholerae with increased ATPase activity. Structure. 2013;21(9):1707–17. doi: 10.1016/j.str.2013.06.027. PubMed PMID: 23954505; PubMed Central PMCID: PMCPMC3775503.

20. Reindl S, Ghosh A, Williams GJ, Lassak K, Neiner T, Henche AL, et al. Insights into FlaI functions in archaeal motor assembly and motility from structures, conformations, and genetics. Mol Cell. 2013;49(6):1069–82. doi: 10.1016/j.molcel.2013.01.014. PubMed PMID: 23416110; PubMed Central PMCID: PMCPMC3615136.

21. Gai D, Zhao R, Li D, Finkielstein CV, Chen XS. Mechanisms of conformational change for a replicative hexameric helicase of SV40 large tumor antigen. Cell. 2004;119(1):47–60. doi: 10.1016/j.cell.2004.09.017. PubMed PMID: 15454080.

22. Rule CS, Patrick M, Camberg JL, Maricic N, Hol WG, Sandkvist M. Zinc coordination is essential for the function and activity of the type II secretion ATPase EpsE. Microbiologyopen. 2016. doi: 10.1002/mbo3.376. PubMed PMID: 27168165.

23. Salzer R, Herzberg M, Nies DH, Joos F, Rathmann B, Thielmann Y, et al. Zinc and ATP binding of the hexameric AAA-ATPase PilF from Thermus thermophilus: role in complex stability, piliation, adhesion, twitching motility, and natural transformation. J Biol Chem. 2014;289(44):30343–54. doi: 10.1074/jbc.M114.598656. PubMed PMID: 25202014; PubMed Central PMCID: PMCPMC4215219.

24. Robien MA, Krumm BE, Sandkvist M, Hol WG. Crystal structure of the extracellular protein secretion NTPase EpsE of Vibrio cholerae. J Mol Biol. 2003;333(3):657–74. PubMed PMID: 14556751.

25. Girdlestone C, Hayward S. The DynDom3D Webserver for the Analysis of Domain Movements in Multimeric Proteins. J Comput Biol. 2016;23(1):21–6. doi: 10.1089/cmb.2015.0143. PubMed PMID: 26540459.

26. Mancl JM, Black WP, Robinson H, Yang Z, Schubot FD. Crystal Structure of a Type IV Pilus Assembly ATPase: Insights into the Molecular Mechanism of PilB from Thermus thermophilus. Structure. 2016. doi: 10.1016/j.str.2016.08.010. PubMed PMID: 27667690.

27. Karuppiah V, Hassan D, Saleem M, Derrick JP. Structure and oligomerization of the PilC type IV pilus biogenesis protein from Thermus thermophilus. Proteins. 2010;78(9):2049–57. doi: 10.1002/prot.22720. PubMed PMID: 20455262.

28. McCallum M, Tammam S, Little DJ, Robinson H, Koo J, Shah M, et al. PilN Binding Modulates the Structure and Binding Partners of the Pseudomonas aeruginosa Type IVa Pilus Protein PilM. J Biol Chem. 2016;291(21):11003–15. doi: 10.1074/jbc.M116.718353. PubMed PMID: 27022027.

29. Georgiadou M, Castagnini M, Karimova G, Ladant D, Pelicic V. Large-scale study of the interactions between proteins involved in type IV pilus biology in Neisseria meningitidis: characterization of a subcomplex involved in pilus assembly. Mol Microbiol. 2012;84(5):857–73. doi: 10.1111/j.1365-2958.2012.08062.x. PubMed PMID: 22486968.

30. Nivaskumar M, Bouvier G, Campos M, Nadeau N, Yu X, Egelman EH, et al. Distinct docking and stabilization steps of the Pseudopilus conformational transition path suggest rotational assembly of type IV pilus-like fibers. Structure. 2014;22(5):685–96. doi: 10.1016/j.str.2014.03.001. PubMed PMID: 24685147; PubMed Central PMCID: PMCPMC4016124.

31. Wang YA, Yu X, Ng SY, Jarrell KF, Egelman EH. The structure of an archaeal pilus. J Mol Biol. 2008;381(2):456–66. doi: 10.1016/j.jmb.2008.06.017. PubMed PMID: 18602118; PubMed Central PMCID: PMCPMC2570433.

32. Craig L, Volkmann N, Arvai AS, Pique ME, Yeager M, Egelman EH, et al. Type IV pilus structure by cryo-electron microscopy and crystallography: implications for pilus assembly and functions. Mol Cell. 2006;23(5):651–62. doi: 10.1016/j.molcel.2006.07.004. PubMed PMID: 16949362.

33. Craig L, Taylor RK, Pique ME, Adair BD, Arvai AS, Singh M, et al. Type IV pilin structure and assembly: X-ray and EM analyses of Vibrio cholerae toxin-coregulated pilus and Pseudomonas aeruginosa PAK pilin. Mol Cell. 2003;11(5):1139–50. PubMed PMID: 12769840.

34. Kolappan S, Coureuil M, Yu X, Nassif X, Egelman EH, Craig L. Structure of the Neisseria meningitidis Type IV pilus. Nat Commun. 2016;7:13015. doi: 10.1038/ncomms13015. PubMed PMID: 27698424.

35. Banerjee A, Neiner T, Tripp P, Albers SV. Insights into subunit interactions in the Sulfolobus acidocaldarius archaellum cytoplasmic complex. FEBS J. 2013;280(23):6141–9. doi: 10.1111/febs.12534. PubMed PMID: 24103130.

36. Camberg JL, Johnson TL, Patrick M, Abendroth J, Hol WG, Sandkvist M. Synergistic stimulation of EpsE ATP hydrolysis by EpsL and acidic phospholipids. EMBO J. 2007;26(1):19–27. doi: 10.1038/sj.emboj.7601481. PubMed PMID: 17159897; PubMed Central PMCID: PMC1782372.

37. Sakes A, van der Wiel M, Henselmans PW, van Leeuwen JL, Dodou D, Breedveld P. Shooting Mechanisms in Nature: A Systematic Review. PLoS One. 2016;11(7):e0158277. doi: 10.1371/journal.pone.0158277. PubMed PMID: 27454125; PubMed Central PMCID: PMCPMC4959704.

38. Patek SN, Korff WL, Caldwell RL. Biomechanics: deadly strike mechanism of a mantis shrimp. Nature. 2004;428(6985):819–20. doi: 10.1038/428819a. PubMed PMID: 15103366.

39. Kabsch W. Xds. Acta Crystallogr D Biol Crystallogr. 2010;66(Pt 2):125–32. doi: 10.1107/S0907444909047337. PubMed PMID: 20124692; PubMed Central PMCID: PMCPMC2815665.

40. Strong M, Sawaya MR, Wang S, Phillips M, Cascio D, Eisenberg D. Toward the structural genomics of complexes: crystal structure of a PE/PPE protein complex from Mycobacterium tuberculosis. Proc Natl Acad Sci U S A. 2006;103(21):8060–5. doi: 10.1073/pnas.0602606103. PubMed PMID: 16690741; PubMed Central PMCID: PMCPMC1472429.

41. Adams PD, Afonine PV, Bunkoczi G, Chen VB, Davis IW, Echols N, et al. PHENIX: a comprehensive Python-based system for macromolecular structure solution. Acta Crystallogr D Biol Crystallogr. 2010;66(Pt 2):213–21. doi: 10.1107/S0907444909052925. PubMed PMID: 20124702; PubMed Central PMCID: PMC2815670.

42. Winn MD, Ballard CC, Cowtan KD, Dodson EJ, Emsley P, Evans PR, et al. Overview of the CCP4 suite and current developments. Acta Crystallogr D Biol Crystallogr. 2011;67(Pt 4):235–42. doi: 10.1107/S0907444910045749. PubMed PMID: 21460441; PubMed Central PMCID: PMCPMC3069738.

43. Emsley P, Lohkamp B, Scott WG, Cowtan K. Features and development of Coot. Acta Crystallogr D Biol Crystallogr. 2010;66(Pt 4):486–501. doi: 10.1107/S0907444910007493. PubMed PMID: 20383002; PubMed Central PMCID: PMC2852313.

44. Afonine PV, Grosse-Kunstleve RW, Echols N, Headd JJ, Moriarty NW, Mustyakimov M, et al. Towards automated crystallographic structure refinement with phenix.refine. Acta Crystallogr D Biol Crystallogr. 2012;68(Pt 4):352–67. doi: 10.1107/S0907444912001308. PubMed PMID: 22505256; PubMed Central PMCID: PMCPMC3322595.

45. Finn RD, Clements J, Arndt W, Miller BL, Wheeler TJ, Schreiber F, et al. HMMER web server: 2015 update. Nucleic Acids Res. 2015;43(W1):W30–8. doi: 10.1093/nar/gkv397. PubMed PMID: 25943547; PubMed Central PMCID: PMCPMC4489315.

46. Sievers F, Wilm A, Dineen D, Gibson TJ, Karplus K, Li W, et al. Fast, scalable generation of high-quality protein multiple sequence alignments using Clustal Omega. Mol Syst Biol. 2011;7:539. doi: 10.1038/msb.2011.75. PubMed PMID: 21988835; PubMed Central PMCID: PMC3261699.

47. Tamura K, Peterson D, Peterson N, Stecher G, Nei M, Kumar S. MEGA5: molecular evolutionary genetics analysis using maximum likelihood, evolutionary distance, and maximum parsimony methods. Mol Biol Evol. 2011;28(10):2731–9. doi: 10.1093/molbev/msr121. PubMed PMID: 21546353; PubMed Central PMCID: PMC3203626.

48. Crooks GE, Hon G, Chandonia JM, Brenner SE. WebLogo: a sequence logo generator. Genome Res. 2004;14(6):1188–90. doi: 10.1101/gr.849004. PubMed PMID: 15173120; PubMed Central PMCID: PMCPMC419797.

49. Karplus PA, Diederichs K. Linking crystallographic model and data quality. Science. 2012;336(6084):1030–3. doi: 10.1126/science.1218231. PubMed PMID: 22628654; PubMed Central PMCID: PMC3457925.

50. Chen VB, Arendall WB, 3rd, Headd JJ, Keedy DA, Immormino RM, Kapral GJ, et al. MolProbity: all-atom structure validation for macromolecular crystallography. Acta Crystallogr D Biol Crystallogr. 2010;66(Pt 1):12–21. doi: 10.1107/S0907444909042073. PubMed PMID: 20057044; PubMed Central PMCID: PMC2803126.

51. Ashkenazy H, Abadi S, Martz E, Chay O, Mayrose I, Pupko T, et al. ConSurf 2016: an improved methodology to estimate and visualize evolutionary conservation in macromolecules. Nucleic Acids Res. 2016. doi: 10.1093/nar/gkw408. PubMed PMID: 27166375.

52. Afonine PV, Moriarty NW, Mustyakimov M, Sobolev OV, Terwilliger TC, Turk D, et al. FEM: feature-enhanced map. Acta Crystallogr D Biol Crystallogr. 2015;71(Pt 3):646–66. doi: 10.1107/S1399004714028132. PubMed PMID: 25760612; PubMed Central PMCID: PMCPMC4356370.

